# Biocatalytic quantification of α-glucan in particulate marine organic matter

**DOI:** 10.1101/2021.11.10.468175

**Authors:** Nicola Steinke, Silvia Vidal-Melgosa, Mikkel Schultz-Johansen, Jan-Hendrik Hehemann

**Affiliations:** University of Bremen, MARUM - Center for Marine Environmental Sciences, Faculty of Biology and Chemistry, Leobener Straße 8, 28359 Bremen, Germany; Max Planck Institute for Marine Microbiology, Celsiusstraße 1, 28359 Bremen, Germany

## Abstract

Marine algae drive the marine carbon cycle, converting carbon dioxide into organic material. A major component of this produced biomass is a variety of glycans; and yet their chemical composition and individual involvement in production, sedimentation and bacterial uptake remain largely unknown due to a lack of analytical tools for glycan-specific quantification.

Marine α-glucans include a range of storage glycans from red and green algae, bacteria, fungi and animals. Although these compounds are likely to account for a high amount of the carbon stored in the oceans they have not been quantified in marine samples so far.

Here we present a method to extract and quantify α-glucans in particulate organic matter from algal cultures and environmental samples using a sequential physicochemical extraction and enzymes as α-glucan-specific probes. This enzymatic assay is more specific and less susceptible to side reactions than chemical hydrolysis. Using HPAEC-PAD to detect the hydrolysis products allows for a glycan quantification in particulate marine samples even at low concentration of ≈ 2-7 µg/L α-glucans.

We measured α-glucans (and compared their concentration with the β-glucan laminarin) in three microalgae laboratory cultures as well as in marine particulate organic matter from the North Sea and western North Atlantic Ocean. While laminarin from diatoms and brown algae is an essential component of marine carbon turnover, our results further indicate the significant contribution of starch-like α-glucans to marine particulate organic matter.

Henceforth, the combination of glycan-linkage-specific enzymes and chromatographic hydrolysis product detection can provide a powerful tool in the exploration of marine glycans and their role in the global carbon cycle.

## 1. Introduction

Autotrophic organisms in the oceans are estimated to contribute about 46 % of the global primary production, converting carbon dioxide to organic compounds (Field et al., 1998). Major components of this biomass are glycans (Myklestad, 1974). While most of the marine glycans are consumed and eventually reconverted into carbon dioxide, marine glycans can also be a carbon sink and become part of the ocean floor sediment. This carbon storage capacity makes marine glycans an important subject for environmental - and particularly climate-related research (Engel et al., 2004; Hedges et al., 2001).

Glycans are complex and often non-linear polymers built from numerous possible monosaccharides (sometimes with chemical modifications) and linkage possibilities (Laine, 1994). This complexity makes the structural analysis of glycans challenging and the analysis of glycans in marine dissolved or particulate organic matter even more difficult as the glycans are present in complex mixtures and at often low individual concentrations. The lack of structure-specific analytical tools for marine glycan samples leads to these glycans being commonly identified and quantified by their monosaccharide content after chemical hydrolysis (Engel and Händel, 2011; Panagiotopoulos and Sempéré, 2005). This non-selective acid hydrolysis destroys information about glycosidic linkages and some chemical modifications. It is therefore not suitable to unequivocally identify different types of glycans. This results in a fundamental lack of knowledge about the structures and functions of marine polysaccharides in the marine carbon cycle (Hedges et al., 2001).

Glucans are glycans derived from D-glucose residues. Some glucans like starch, glycogen and laminarin are energy storage glycans while other glucans like cellulose are structural polymers (Suzuki and Suzuki, 2013; Painter, 1983). Acid hydrolysis makes these glycans of different biological origins indistinguishable and obscures insight into their roles in the marine carbon cycle.

α-Glucans include a range of different types of polysaccharides, most of them containing α-1,4 and α-1,6 glycosidic linkages. Starch is a mixture of the linear α-1,4-linked amylose and the branched amylopectin composed of α-1,4- and α-1,6-glycosidic bonds (Imberty et al., 1991). Starch is commonly known as a storage glycan for terrestrial plants but it is also found in green algae. Floridean starch is a storage α-glucan in red algae similar to amylopectin but with a higher degree of α-1,6-branching. Glycogen is another highly α-1,6-branched storage α-glucan found in animals, fungi and bacteria (Ball and Morell, 2003). Although the amount of α-glucans in the ocean is unknown, analysis of marine bacterial polysaccharide degradation pathways (Fang et al., 2019; Kappelmann et al., 2019) indicate that α-glucans are an important carbohydrate source for marine bacteria.

Glycan specific glycoside hydrolases (GHs) can be applied to identify and quantify α-glucans in the ocean. There are several well-characterized starch specific GHs (α-amylases and amyloglucosidases) that have been used to quantify starch in food (McCleary et al., 1994, 2002), yet a more sensitive method for starch quantification in marine environmental samples is missing.

In this study, a set of commercial enzymes was adapted into an assay for quantification of α-glucans in different types of microalgae and unprocessed marine environmental samples. This assay was tested in parallel with a previously developed laminarin assay (Becker et al., 2020) on different types of α-glucans and other polysaccharides to explore possible side reactions and compare the enzymatic hydrolysis (EH) with the commonly used acid hydrolysis (AH). Additionally, the detection range of the assay was tested using two types of glucose detection, the spectrophotometric PAHBAH assay and the chromatographic HPAEC-PAD. Furthermore, the extraction of glucans from different types of microalgae was optimized. The α-glucan and laminarin content of these microalgae was tested at different days of algal growth and compared to the total particulate organic carbon (POC) in these samples. Finally, the assay was used on two sets of particulate organic matter (POM) samples from the North Sea (spring 2020) and the western North Atlantic Ocean (spring 2019) to demonstrate that this method allows for a quantification of low concentrations of α-glucans alongside laminarin in crude marine samples.

## 2. Methods

### 2.1. Algal cultures

Three species of microalgae were used as examples for marine organic material to test different extraction protocols and enzymatic hydrolysis of algal glucans: The green microalga *Ostreococcus tauri* (Ral et al., 2004) was grown in L1 medium (Guillard, 1983; Guillard and Hargraves, 1993), and the red microalgae *Porphyridium purpureum* and the diatom *Thalassiosira weissflogii* were grown in ESAW medium (Harrison et al., 1980). Algae cultures were kept at a constant temperature of 15°C, with a 12-h/12-h light/dark cycle, without stirring and irradiated with *≈* 140 µmol photons m^−2^s^−1^.

For glucan extraction tests, duplicate T75 flasks with 400 mL growth medium were inoculated with 5 mL of 7-days old algal cultures. After 20 days the material was filtered at 200 mbar on a combusted (450 °C, 4 h) 25 mm GF/F glass microfiber filter (using 20 mL of algal culture per filter). The filters were stored at -20°C until extraction.

For the α- and β-glucan quantifications algae were cultivated in triplicate batch cultures. A total volume of 250 mL in T75 suspension cell culture flasks was inoculated with 5 mL of a 25-mL culture that had been grown for 7 days. 10-25 mL of each culture was taken over 20 days and filtered as described above.

### 2.2. Environmental sample collection

Sampling in the North Atlantic Ocean (40°53.7’N, 60°11.9’W) was carried out in May 2019 and in the North Sea (54°11.3’N, 7°54.0’E) in April 2020. In both locations, surface seawater was collected and directly filtered through precombusted (400 °C, 4 h) GF/F glass filters with 142 mm diameter (Whatman glass microfiber filters, WHA1825142, Sigma-Aldrich). For the Atlantic Ocean samples, 50 L seawater were filtered through each GF/F filter with a peristaltic pump (Watson Marlow 630 S) at 40 rpm. Filters were stored at -80 °C until further analysis. For the North Sea samples, 15 L seawater were filtered through each GF/F filter with an air pressure pump (Flojet G57, ITT Industries) at *≈* 0.2-0.5 bar. Filters were stored at -30 °C until further analysis.

### 2.3. Extraction of glucans from particulate algae material

The filters were cut into equally sized pieces and subjected to extraction tests. One filter piece was kept as a non-extracted reference. Each extraction was tested using filter-part triplicates of three different filters. Afterwards the extracted and non-extracted filter-parts were put under AH conditions and the glucose content of the acid extract tested using HPAEC-PAD. The extraction conditions tested include water or 1M NaOH extractions at 99°C or in a sonication bath (SONOREX SUPER RK 510, Bandelin), different extraction times and sequential extractions.

### 2.4. Acid hydrolysis

Glycan samples in solution or algal samples on glass fiber filters were incubated in 1 M HCl for 24 h at 100°C in sealed glass ampoules to chemically hydrolyze glycans. Afterwards, 100 µL of each acid solution was evaporated using a speed-vac (RVC 2-18 CDplus HCl resistant, Christ) and resuspended in 100 µL buffer or Milli-Q water.

### 2.5. Enzymatic hydrolysis

α-Glucans were hydrolyzed using α-amylase (*Aspergillus oryzae*) obtained from Megazyme (Product code: E-ANAAM) and amyloglucosidase (*Aspergillus niger*) from Merck (Product code: 10115). The stock concentration was 1 U/µL for both enzymes. Samples containing polysaccharide standards or extracted glucans were split into six subsamples: three for enzyme hydrolysis and three for non-hydrolyzed controls.

Triplicate subsamples of 90 µL aqueous glycan extract were mixed with 10 µL sodium acetate buffer (1 M, pH 4.5), 0.4 µL α-amylase stock (in 3.2 M ammonium sulphate), 0.9 µL amyloglucosidase stock (in 0.1 M sodium acetate) and 1 µL BSA solution (100 mg/mL). Triplicate non-hydrolyzed subsamples were prepared in the same way, but contained 0.4 µL 3.2 M ammonium sulphate and 0.9 µL 0.1 M sodium acetate instead of enzymes. Reaction contents were mixed and incubated for 35 min at 50 °C (and 400 rpm shaking) in a heat block. Then, the enzyme reactions were inactivated 5 min at 99 °C, centrifuged (10.000 rpm for 30 s) and cooled on ice.

Subsamples used for laminarin quantification (90 µL) were hydrolyzed using 90 µL aqueous glycan solution, 10 µL 500 mM MOPS buffer (pH 7.0) and laminarinases as previously described (Becker et al., 2017).

### 2.6. PAHBAH reducing sugar assay

Photometric quantification of polysaccharide hydrolysis products was performed using the PAHBAH reducing sugar assay (Lever, 1972). One milliliter of a freshly prepared 9:1 (v/v) mixture of PAHBAH reagent A (0.3 M 4-hydroxybenz-hydrazide, 0.6 M HCl) and PAHBAH reagent B (48 mM trisodium citrate, 10 mM CaCl_2_, 0.5 M NaOH) was added to 0.1 mL of sample. After incubation for 5 min at 99 °C, the samples were cooled on ice and the absorbance at 410 nm was measured using 10 mm pathlength semimicro cuvets and a BioSpectrometer (Eppendorf). Glucose standards were prepared and measured in the same manner.

### 2.7. HPAEC–PAD

All samples for HPAEC–PAD (High-performance Anion Exchange Chromatography with pulsed amperometric detection) were filtered using a 0.2 µm Spin-X filter and transferred into an HPLC-vial with a micro-insert. The monosaccharide (and specifically glucose) content was determined using HPAEC-PAD with a Dionex CarboPac PA10 column (Thermo Scientific) and monosaccharide mixes as standards for calibration (Engel and Händel, 2011). The oligosaccharide content was determined using Dionex CarboPac PA200 and PA100 columns (Thermo Scientific) and appropriate malto- and laminarin oligosaccharides (Megazyme).

### 2.8. Glucan quantification

Glucans in extracted samples were quantified as glucose equivalents based on a calibration curve generated with glucose standards.

The glucose signals were inferred from PAHBAH absorbance readings or integrated peak areas from HPAEC-PAD chromatograms. Each hydrolyzed sample or non-hydrolyzed control was measured in triplicates. To correct for background signals in the enzymatically digested samples, the glucose values detected for non-hydrolyzed samples were subtracted. No correction was applied for total glucan quantification after acid hydrolysis.

The glucan concentrations in filter extracts were normalized by extraction volume, filter size and filtration volume.

### 2.9. Particulate organic carbon quantification

Particulate organic matter from microalgae cultures or environmental samples were both filtered on glass fiber filters. Defined pieces of these filters were punched out in triplicates and subjected to an acidic atmosphere with concentrated HCl for 24 h in a desiccator to remove inorganic carbon. Afterwards, the filters were dried for 24 h at 60 °C and wrapped in combusted tin foil. The carbon quantification was performed by an elemental analyzer (vario MICRO cube; Elementar Analysensysteme) using sulfanilamide as a calibration standard.

## 3. Assessment

### 3.1. Optimal conditions for enzymatic *α*-glucan hydrolysis

Based on an assay for amylose and amypectin quantification in cereal starches and flours (Megazyme, 2018), a similar assay for total α-glucan quantification (without amylose/amylopectin separation) was developed and optimized for a sample volume of 100 µL, a starch concentration of 100 µg/mL (from corn, Sigma-Aldrich) and hydrolysis product detection using the PAHBAH reducing sugar assay (Lever, 1972). Optimal EH was achieved by incubating a 100 µL starch sample in 100 mM sodium acetate buffer (pH 4.5) for 35 min at 50 °C in the presence of 0.4 U of the *endo*-acting α-amylase (from *Aspergillius oryzae*, Megazyme) and 0.9 U of the *exo*-acting amyloglucosidase (from *Aspergillus niger*, Merck). Under these conditions, PAHBAH reducing sugar signals were maximal (Figures S1, S2). Subsequently, the complete hydrolysis of different α-glucan polysaccharides, namely starch, glycogen, amylose or amylopectin down to glucose monosaccharides was confirmed by HPAEC-PAD (Figure S3).

### 3.2. Enzymatic hydrolysis of *α*-glucans is more effective than acid hydrolysis

The enzymatic α-glucan hydrolysis was tested on different starch concentrations and compared to AH using the PAHBAH reducing sugar assay as a glucose detection method (Figure 1). The limit of quantification was found to be *≈* 11 µg/mL starch, both for acid hydrolysis, and for enzymatic hydrolysis in combination with the PAHBAH assay. However, it was found that the EH of starch (1 1000 µg/mL) generated a higher photometric signal than AH suggesting that a more complete polysaccharide hydrolysis was achieved or that EH is less prone to generate side products (Cai et al., 2014). Especially at higher starch concentrations above 250 µg/mL the AH glucose signal visibly flattened, resulting in a nonlinear regression line. In contrast, EH of starch followed a linear regression model for all measured starch concentrations up to 1 mg/mL.

**Figure 1:**
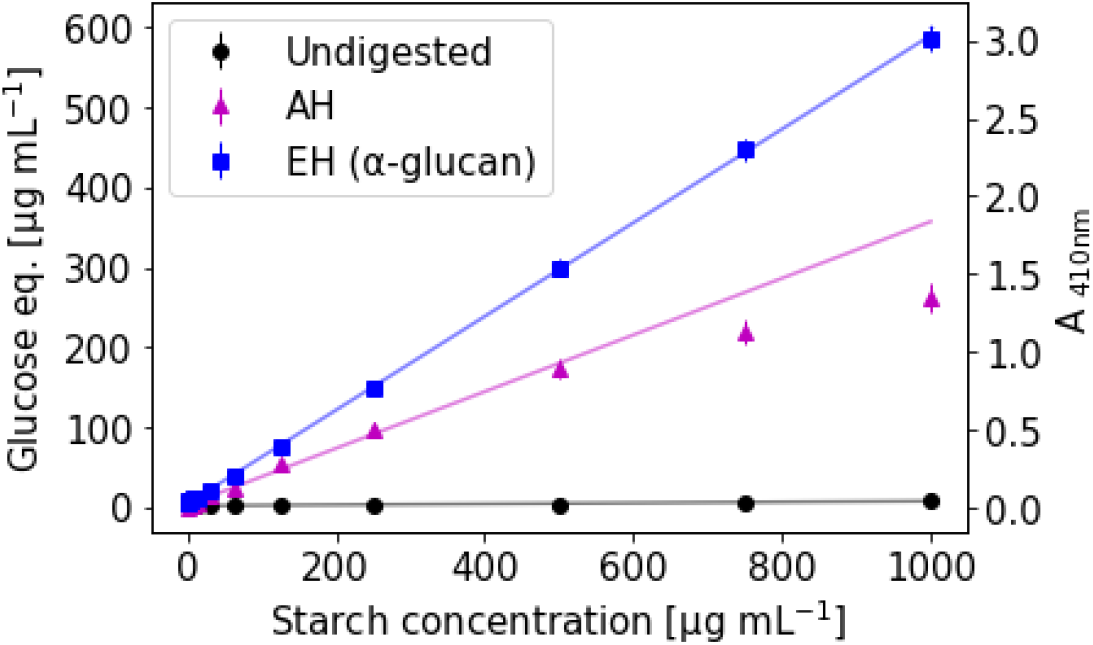
Enzymatic hydrolysis is more effective than acid hydrolysis for starch: Enzymatic hydrolysis (EH) products of starch, acid hydrolysis (AH) products of starch, non-digested starch, and glucose as calibration standard were measured in α-glucan assay buffer (100 mM sodium acetate buffer, pH 4.5) using the PAHBAH reducing sugar assay. A_410nm_ signals above 1.9 were measured in diluted samples. Error bars denote standard deviation (n=3, technical replicates). Solid lines represent regression lines.

A complete hydrolysis of 1 g of a glucan into glucose would theoretically yield approximately 1.1 g of glucose by the addition of 1 water molecule per hydrolyzed glucose molecule. In comparison, Figure 1 shows that even EH is not complete, as 1 mg/mL starch yield 0.6 mg/mL of glucose equivalents. This result is probably due to the water insolubility of some starch particles, and because the amount of enzymes added was optimized for 100 µg/mL starch samples.

It should be noted that for these measurements, the glucose calibration curve was prepared in water. Under AH conditions the glucose signals in the calibration curve were (*≈* -20 %) lower (Figure S4). These data indicate that lower glucose signals in acid hydrolyzed starch samples could partially be due to acid catalyzed conversion of glucose. But as this loss does not account for the total difference in signal of acid and enzymatic starch hydrolysis, our results also indicate that AH of starch is less complete than EH.

Overall, these results show that AH - the traditional quantification method may underestimate the true glycan content in environmental samples due to incomplete glycan hydrolysis and possible side reactions reducing the detectable monosaccharide content.

### 3.3. Enzymatic hydrolysis of *α*-glucans is specific

To explore the specificity of the enzymatic α-glucan assay compared with AH, 14 commercially available polysaccharides were tested. The enzymatic laminarin assay was included as a control. Figure 2 shows the reducing sugar assay results for the 8 polysaccharides that could be hydrolyzed by either of the enzymatic assays compared to AH. Additional tested polysaccharides that could only be hydrolyzed by acid are listed in Table S1.

**Figure 2:**
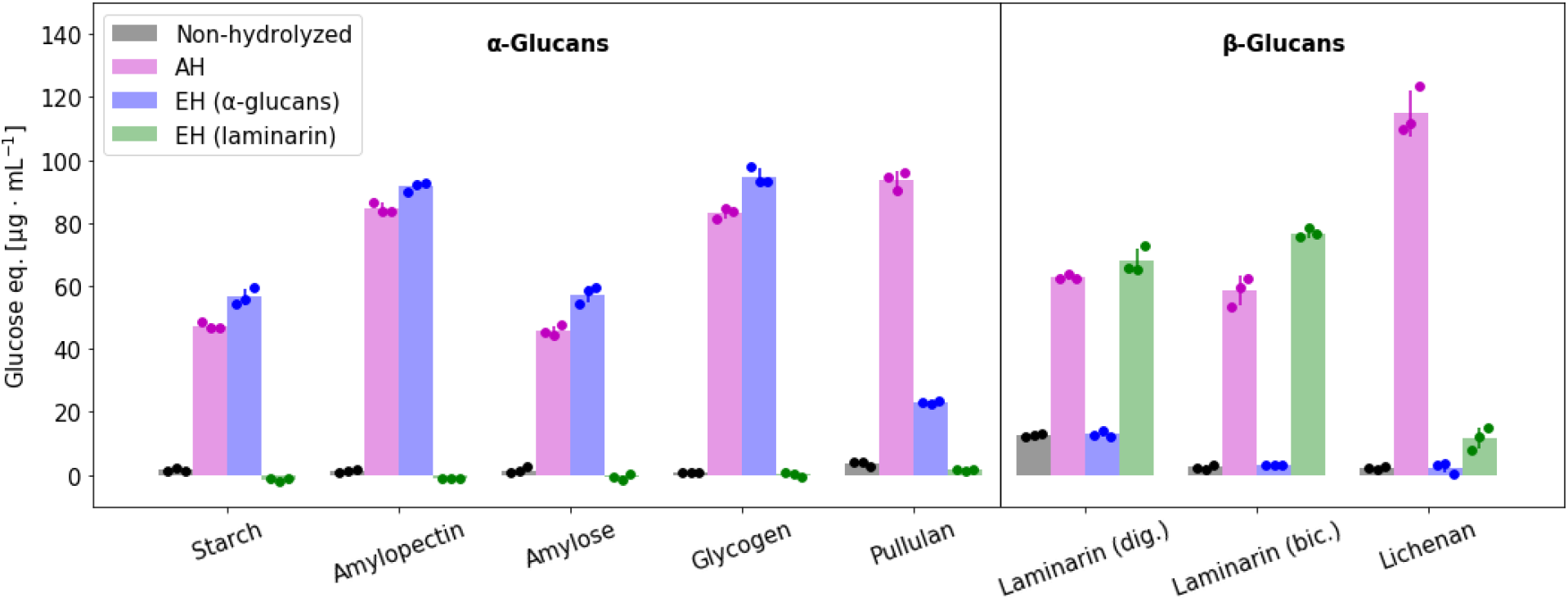
Enzymatic hydrolysis of α-glucans is specific and more effective than acid hydrolysis: Hydrolysis efficiency of different polysaccharides (100 µg/mL) using acid hydrolysis (AH) compared to enzymatic hydrolysis (EH) using the α-glucan or laminarin assay. All tested polysaccharides are listed in Table S1. Only the results of polysaccharides that could be hydrolyzed by either of the enzymatic assays are depicted. Data points represent individual samples, and error bars denote standard deviation (n=3, technical replicates).

To compare the measured glucans, glucose was used as a quantification standard. The glucose standards used for quantification were treated the same way as the respective samples. This corrects for absorbance changes in the assay caused by addition of specific buffers and enzymes, or by AH (Figure S4).

The α-glucan assay is able to hydrolyze all polysaccharides containing α-glucan 1-4- and 1-6-linkages, namely amylopectin, amylose, glycogen and pullulan. The reducing sugar signal of enzymatically hydrolyzed amylopectin and glycogen equals approximately 100 µg/mL glucose indicating a complete hydrolysis to glucose. Amylose is poorly soluble in water and therefore only 60 µg/mL glucose equivalents, presumably produced by shorter-chain amylose, could be measured using EH.

Similar to the results in Figure 1 the concentration of glucose hydrolysis products is often higher for the α-glucan enzymatic assay than for the AH with the notable exception of pullulan, which is less effectively hydrolyzed by the enzymatic assay than by AH. Pullulan is comprised of α-1-4-maltotriose units linked by α-1-6-linkages and should - in theory - be completely hydrolyzed by the applied *endo*-1-4-α-amylase and *exo*-1,4/1,6-α-amyloglucosidase. However, as the observed reducing sugar signal is only slightly increased by the application of these enzymes, it can be assumed that this α-amylase needs a longer uninterrupted α-1-4-linked glucose chain to be more active. Thus, the α-glucan assay can mainly be used to identity and quantify starch-like α-glucans with 1-4 chains and optional 1-6 branches.

While the laminarin assay is effective on both laminarins tested and the amount of released glucose is similar and slightly higher than than for AH, the only other polysaccharide being partially hydrolyzed by this assay is lichenan. As lichenan is comprised of β-1-3 and β-1-4 linked glucose units and some of the applied laminarinases have a β-1-3 activity (Becker et al., 2017), this small side reaction is to be expected. However, the results show that the laminarin assay cannot detect β-glucan from barley and any of the tested α-glucans.

The relatively high reducing sugar signal of non-hydrolyzed laminarin from *Laminaria digitata* indicates the presence of shorter polysaccharide chains. In contrast, non-hydrolyzed polysaccharide from other batches of the same laminarin had no measureable reducing sugar signal (Becker et al., 2020). This further underlines the advantage in measuring glycans in glucose equivalents over using commercial polysaccharides as standards.

Overall, these results demonstrate that a combination of the enzymatic assays can distinguish the important storage glucans; starch-like α-1,4/1,6-glucans and laminarin - something that cannot be achieved using traditional chemical hydrolysis.

### 3.4. Microalgal glucans can be extracted using hot water and sonication

The use of enzymatic assays to quantify glucans in marine POM requires that the glucans can be extracted from environmental samples. Particulate microalgal material was used to test different extraction protocols. The extraction efficiency was determined by measuring the ratio of glucose detected by HPAEC-PAD on extracted and non-extracted filter pieces after AH conditions, assuming that these conditions are sufficient for a complete glucan extraction and hydrolysis. For the extraction of glucans from filters, we tested different combinations of water and NaOH incubations with, and without, sonication.

Figure 3 shows the results of these extraction condition tests as the proportion of (total) glucose that remains in the filters after extraction. Water extraction for 1 h at 99 °C is able to extract more than 90 % of *T. weissflogii* particulate glucans. For *O. tauri*, only up to 50 % of glucans are extracted under the same conditions (Figure 3 A). A longer incubation time, sequential water extractions or a hot alkaline extraction using 1 M sodium hydroxide (1 h, 99 °C) do not increase the extraction efficiency for *O. tauri*. However, a subsequent additional extraction using 1 M sodium hydroxide (1 h, 99 °C) decreases the amount of residual glucose to approximately 30 % (Figure 3 B). Sonication of water extracted samples before incubation at 99 °C gives similar results; approximately 25 % residual glucose for *O. tauri* and 30 % residual glucose for *P. purpureum* (Figure 3 C).

**Figure 3:**
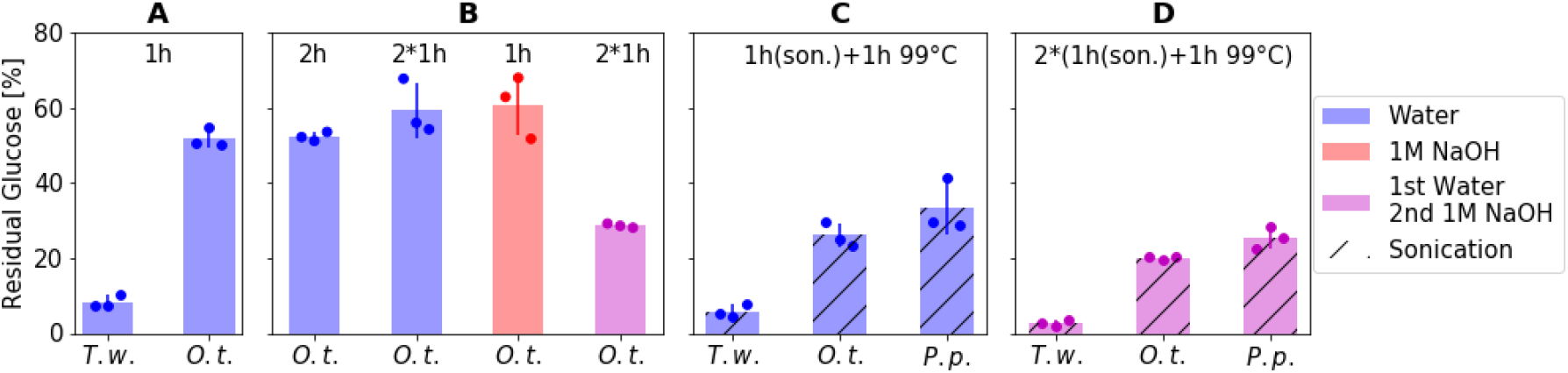
α-Glucans can be extracted from microalgae: Residual glucose content of algal POM on glass fiber filters after extractions (see below) and subsequent AH was determined using HPAEC-PAD. (A) Hot water extraction (1 h, 99 °C) of *T. weissflogii* (*T. w*.) and *O. tauri* (*O. t*.) POM. (B) *O. tauri* extractions with a longer incubation time (water, 2 h, 99 °C), sequential double extraction (water, 2 ×1 h, 99 °C), alkaline extraction (1 M sodium hydroxide, 1h, 99 °C) or sequential water (1 h, 99 °C) and alkaline (1 M sodium hydroxide, 1 h, 99 °C) extraction. (C) Hot water extraction of microalgae (*T. weissflogii, O. tauri, P. purpureum* = *P. p*.) with prior sonication treatment (1 h sonication bath, 1 h, 99 °C), (D) additional subsequent alkaline extraction (in 1 M sodium hydroxide), 1 h sonication bath, 1 h, 99 °C. Individual data points are shown in darker colors, error bars denote standard deviation (n=3, technical replicates).

The extraction was slightly improved when using 1 h sonication in 1 M NaOH followed by 1 h at 99 °C. This treatment resulted in residual glucose values of 3 %, 20 % and 25 % in *T. weissflogii, O. tauri* and *P. purpureum*, respectively(Figure 3 D). However, this additional alkaline extraction did not substantially increase the efficiency of the extraction protocol. Therefore, POM samples were extracted using hot water (1 h, sonication; 1 h, 99 °C) in the following sections. The integrity of laminarin and starch was also tested and confirmed by EH of starch and laminarin standards that had been subjected to extraction conditions compared to non-subjected samples.

These results demonstrate that glucans in *O. tauri* and *P. purpureum* are more resistant to aqueous extraction methods than glucans in *T. weissflogii*. This result is probably due to different glucans present in these species. While material of the diatom *T. weissflogii* should contain laminarin (Becker et al., 2017), the green microalgae *Ostreococcus tauri* and the red microalgae *Porphyridium purpureum* should contain starch and flouridean starch respectively (Sorokina et al., 2011; Sheath et al., 1979). Laminarin is highly soluble in water, whereas α-glucans vary in their water solubilities; branched α-glucans like amylopectin and glycogen have a high solubility and linear amylose has a low solubility. The differences in extraction efficiency should be taken into account with regard to glucan quantifications as the true content might be higher.

### 3.5. *α*-Glucans and laminarin can be quantified in parallel in particulate matter extracts from cultivated microalgae

Glucan quantification in POM from *T. weissflogii, O. tauri* and *P. purpureum* laboratory cultures expressed as glucose equivalents. Laminarin and α-glucans were quantified in microalgal POM extracts using enzymatic hydrolysis and subsequent PAHBAH reducing sugar assay (EH, green and blue symbols). Total glucan was quantified by acid hydrolysis and subsequent PAHBAH reducing sugar assay (AH, magenta symbols). The amount of glucose equivalents in each sample was calculated using glucose calibrations curves.

To investigate the proportion of α- and β-glucans in the aforementioned three microalgae, enzymatic α-glucan and laminarin hydrolysis assays have to be used. Three non-axenic cultures of each of the microalgae *O. tauri, P. purpureum* and *T. weissflogii* were grown in the laboratory and samples from each culture were taken over 20 days and filtered onto glass fiber filters.

Glycans were extracted from these filters using sonication and hot water (as described above) and hydrolyzed by AH or EH. Figure 4 shows the glucose equivalents as determined by the reducing sugar assay over the course of 20 days of algal growth. Laminarin was detectable in *T. weissflogii* diatom cultures after 6 days, but could not be detected in cultures of green microalgae *O. tauri* and red microalgae *P. purpureum* over the 20 days incubation period. Conversely, α-glucans were detected in *O. tauri* and *P. purpureum* from day 4 or 2, respectively. Overall, these findings match the polysaccharide profiles of these three microalgae (Becker et al., 2017; Sorokina et al., 2011; Sheath et al., 1979). However, a small amount of α-glucans could also be measured in two of the three *T. weissflogii* cultures on day 10 and 20. As the algal cultures were non-axenic, it is possible that these α-glucans originate from bacteria.

**Figure 4:**
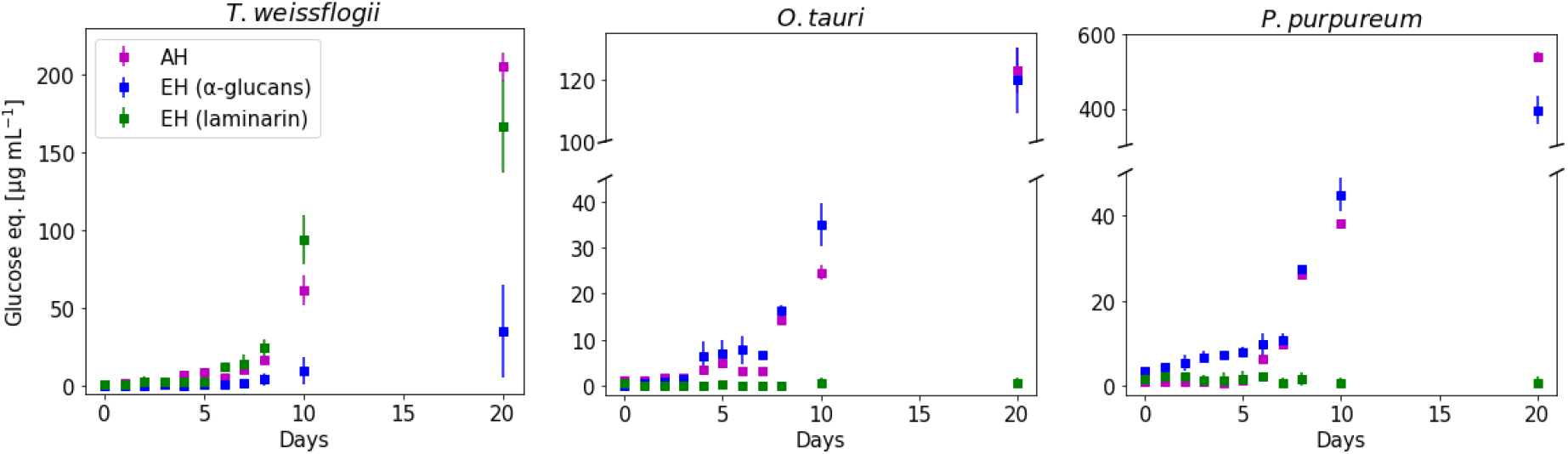
α-Glucans can be quantified alongside laminarin in microalgal extracts using glucan specific enzymatic hydrolysis: Glucan quantification in POM from *T. weissflogii, O. tauri* and *P. purpureum* laboratory cultures expressed as glucose equivalents. Laminarin and α-glucans were quantified in microalgal POM extracts using enzymatic hydrolysis and subsequent PAHBAH reducing sugar assay (EH, green and blue symbols). Total glucan was quantified by acid hydrolysis and subsequent PAHBAH reducing sugar assay (AH, magenta symbols). The amount of glucose equivalents in each sample was calculated using glucose calibrations curves. Error bars denote standard deviation (n=3, biological replicates).

Overall, AH produces a similar total glucose signal as both the enzymatic assays combined. Since the PAHBAH assay would be reactive to all reducing sugars released by AH this indicates that α-glucans and laminarin (a β-glucan) are the predominant glycans in these algal species and under the culture conditions and duration used here.

Figure 5 shows the glucan quantification on day 20 in relation to the total POC. It should be noted that POC was quantified directly on glass fiber filters without prior extraction, whereas for glucan quantification the filters were extracted by sonication and hot water. As this extraction might not be complete, the true glucan (especially α-glucan-) content might be higher than depicted.

**Figure 5:**
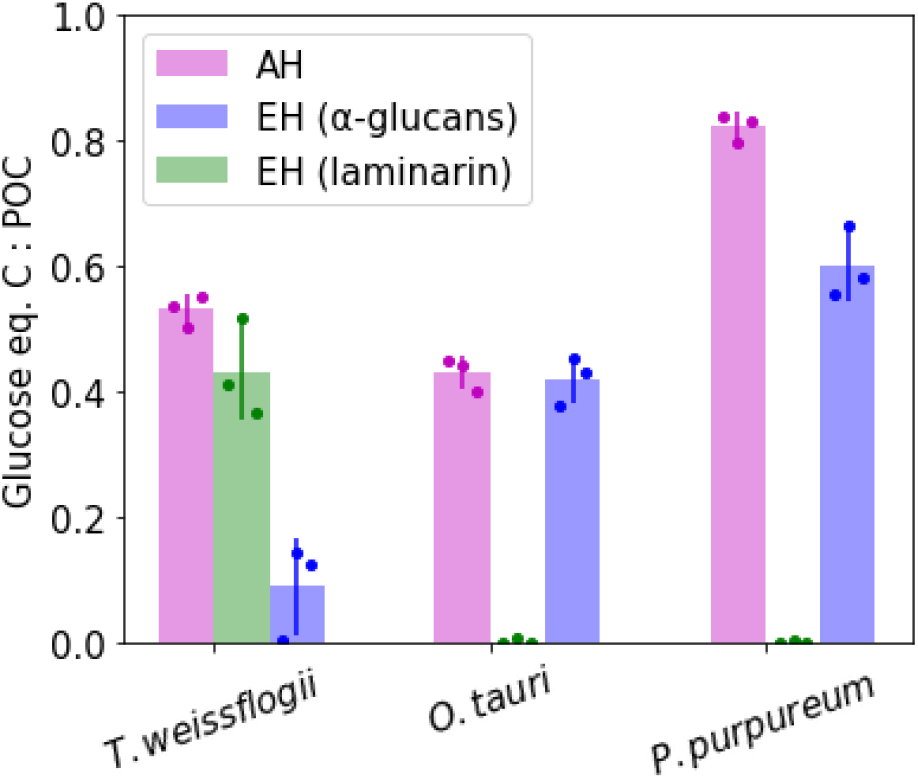
α-Glucans account for a substantial amount of the total POC in red and green algae: α-Glucan and laminarin quantification of microalgal POM from *T. weissflogii, O. tauri* and *P. purpureum* laboratory cultures after 20 days of growth. Glucan quantities are shown as glucose equivalent carbon relative to the total POC. Error bars denote standard deviation (n=3, biological replicates).

For *T. weissflogii* about 40 % of the total POC can be assigned to laminarin. Furthermore, approximately 10 % of POC are detectable in two of three cultures as reducing sugars released by AH. These 10 % are hydrolyzed by enzymatic α-glucan hydrolysis and probably originate from bacterial glycogen. About 40 % of the POC of *O. tauri* corresponds to α-glucans. In this case, the reducing sugars hydrolyzed by acid equal the enzymatically hydrolyzed α-glucans. The POC of *P. purpureum* after 20 days of algal growth can largely be contributed to glycans as approximately 80 % of POC are AH products. However, only about 60 % of *P. purpureum* POC is hydrolyzed by the α-glucan assay. This discrepancy might be due to a higher content of monomeric glucose or other glucose containing oligo- and polysaccharides.

In summary, around 40-60 % of POC in the tested microalgal cultures can be ascribed to glucose coming from glucans. The glucans in *T. weissflogii* are mostly laminarin (β-glucans), the glucans in *O. tauri* and *P. purpureum* are mostly starch-like α-glucans.

### 3.6. *α*-Glucans and laminarin can be quantified in parallel in marine environmental particulate organic matter

Marine environmental POM samples were taken from the western North Atlantic Ocean (40°53.7’N, 60°11.9’W) during May 2019 and in the North Sea (54°11.3’N, 7°54.0’E) near Helgoland (Germany) during April 2020. In both cases, POM in surface water was filtered onto 0.7 µm glass fiber filters. The POC content was quantified directly from non-extracted filter-circle-cutouts, while the glucan content was determined using AH or EH of hot water extracts of quarter filters. The glucose content in the samples was found to be too low for quantification by the photometric reducing sugar assay. Therefore, HPAEC-PAD was used to determine the glucose content in hydrolyzed and non-hydrolyzed extract samples. With HPAEC-PAD, as little as 0.3 µg/mL and 0.2 µg/mL glucose could be detected following AH or EH, respectively, of a starch standard sample. In comparison, the quantification limit for the reducing sugar PAHBAH assay was 11 µg/mL.

HPAEC-PAD analysis of enzymatically hydrolyzed α-glucans and laminarin shows that the only product is glucose (Figure S3). Therefore glucose can be used as a standard to quantify and compare α-glucans and laminarin. Similar chromatograms of hydrolyzed and non-hydrolyzed extracts from algal cultures and environmental samples are depicted in Figure S5.

Figure 6 shows the results of glucan quantification of environmental POM samples (taken at different times over 2-3 days) in relation to the total POC. While laminarin content in POM samples from the western North Atlantic Ocean varies between 2-16 µg/L carbon, it is higher in most of the North Sea samples (4-36 µg/L carbon from laminarin). This is in agreement with the abundance of microalgae at the time of sampling: Samples from the Atlantic Ocean were harvested in the morning (10:30) and evening (22:30) in oligotrophic waters, while North Sea samples were taken in the morning (7:30) and evening (21:30) in May 2020 during the phytoplankton spring bloom. For these North Sea samples a difference between higher α-glucan and laminarin levels in the evening (POC *>* 300 µg/L) and lower levels in the morning (POC *<* 300 µg/L) is observed. This could indicate a diel cycling of α-glucans similar to laminarin during phytoplankton blooms (Becker et al., 2020). This difference in glucan levels depending on time could not be observed in samples from the western North Atlantic Ocean.

**Figure 6:**
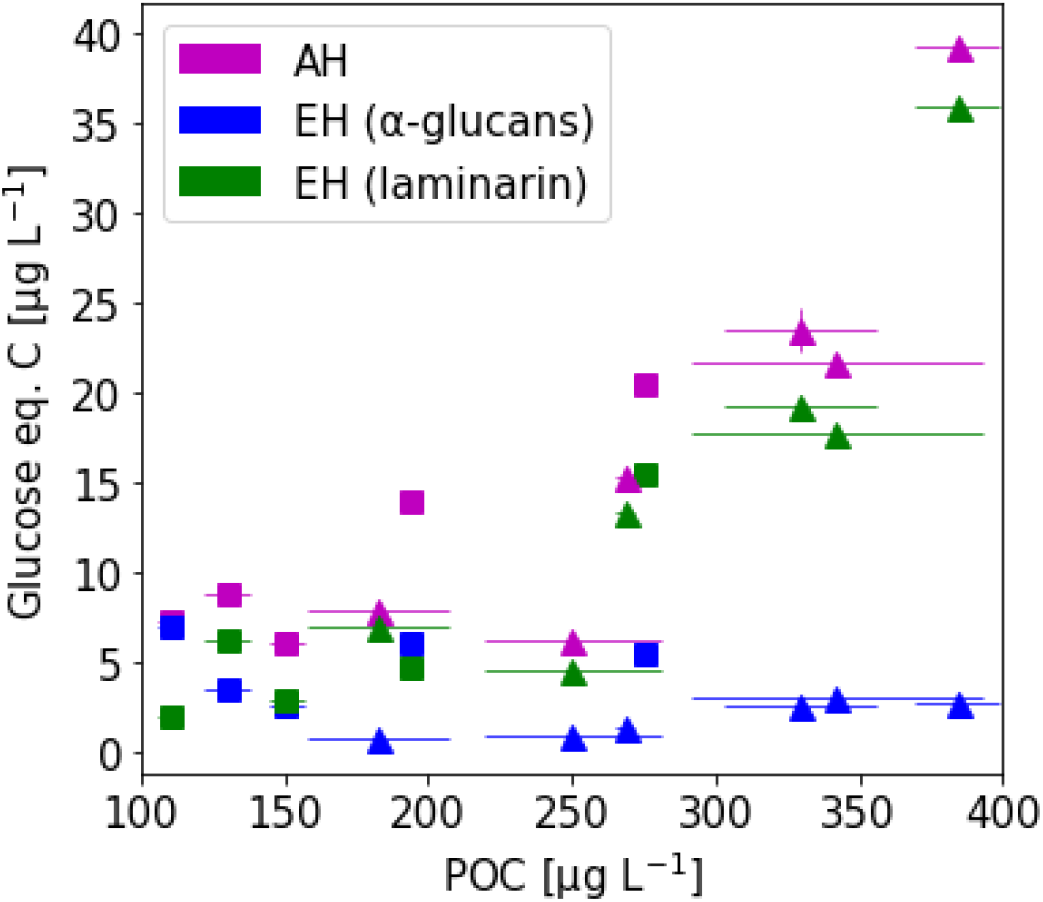
α-Glucans can be quantified alongside laminarin in marine POM samples using glucan specific enzymatic hydrolysis: POM samples on 0.7 µm glass fiber filters were extracted and hydrolyzed using acid hydrolysis (AH) (magenta symbols) or enzymatic hydrolysis (EH) (blue and green symbols). Square symbols represent samples from the western North Atlantic Ocean (May 2019). Triangle symbols represent samples from the North Sea (April 2020). Error bars denote standard deviation (n=3, technical replicates).

Overall, less α-glucan than laminarin was detected in the environmental samples (2-7 µg/L, West Atlantic and 1-4 µg/L, North Sea). The combined glucose value for laminarin and α-glucan, determined with EH, is similar to the total glucose value determined with AH. This observation is consistent with the results shown above (Figures 4 and 5). This result further indicates that most glucose containing polysaccharides are detected by one of the two enzymatic assays. Laminarin content in POM appears to correlate with increasing total POC (*R*^2^ = 77.5 %) - in accordance with previous studies (Becker et al., 2020) -, whereas α-glucan content is significantly lower and apparently non-correlating with total POC (*R*^2^ = 9.4 %). It should be noted that these α-glucans may originate not only from starch-producing red and green algae, but also from glycogen-producing bacteria.

## 4. Discussion

A method for the detection and quantification of starch-like α-glucans in the ocean has been developed and applied to microalgal cultures and marine environment samples. Combined with enzymatic laminarin hydrolysis most of the glucans in marine POM can be quantified.

This method has been shown to be highly specific and robust. However, the assay cannot distinguish between the different types of α-1,4/1,6-glucans and is therefore not suitable to determine the origin of the detected α-glucans. To investigate this question further the degree of α-1,6-branching could also be determined by expanding the assay to include an *endo*-acting α-1,6-isoamylase (Figure S1). Knowing the average degree of α-1,6-branching in POM glucans does not unequivocally identify the type of detected α-glucans, but it might allow for an estimate if the main α-glucan is the highly branched glycogen. Another approach to identify α-glucan types might be the application of separation techniques based on different solubilities.

A related issue requiring further consideration is the hot water (combined with ultrasonication) extraction applied in this study. Varying water solubilities of different α-glucans and the presence of starch granules in algae cells impede a complete glucan extraction from microalgal POM. Previous studies where starch was extracted from algae material also employed ultrasonication (Kobayashi et al., 1974) or other cell disrupting techniques like hot alkaline extraction, bead-beating (Wong et al., 2019) and grinding (Yu et al., 2002). However, for these studies large amounts (kg range) of algae material were used and their aim was rather a high purity of starch extracts and not a complete extraction. Our method of hot water extraction combined with ultrasonication from POM is probably not complete for all types of α-glucans, but it can be employed for small quantities of algae material and it allows parallel detection of other polysaccharides by enzymatic assays. Nevertheless, it is possible that the true α-glucan content of the environmental POM samples is higher than detected.

Furthermore, this work has shed light on problems of glucan quantification after chemical lysis, as this method consistently produced less glucose products from polysaccharide standards than enzymatic hydrolysis. Despite this problem, acid glycan hydrolysis allows for a complete and non-specific hydrolysis and can provide valuable information on monosaccharide composition of environmental samples.

α-Glucan concentrations in marine surface water POM have - in our examples been shown to be lower than laminarin but at a constant level, independent of increased POC concentrations during the seasonal algal bloom in the North Sea 2020. Higher α-glucan concentrations are to be expected near coastal and estuary areas, due to input of terrestrial plants and sites with distinct green and red algal growth. However, it is likely that for marine glycan samples α-glucan concentrations will usually be lower than laminarin concentrations. This low concentration of specific types of glycans in the marine carbon pool is a major obstacle for marine glycan analysis and requires the application of adequate detection methods with low detection limits like HPAEC-PAD or MS.

In conclusion our study has shown that enzymes can be used to identify and quantify glycans in parallel in marine particulate organic matter. The combination of laminarin and α-glucan assays allows for a complete glucan detection in these samples.

## 5. Acknowledgements

This research was supported by the Max Planck Society and supported by the Deutsche Forschungsgemeinschaft (DFG) “Emmy-Noether”-grant HE 7217/1-1, and through the Cluster of Excellence “The Ocean Floor — Earth’s Uncharted Interface” project 390741603 (J.-H.H.).

We thank Carol Arnosti and the captain, crew and scientific party of the RV Endeavor cruise EN638 for helping with sample acquisition in the western North Atlantic Ocean in 2019. This cruise was supported by NSF-OCE1736772 to Carol Arnosti. We thank the AWI-Biologische Anstalt Helgoland team for helping with sample acquisition in the North Sea in 2020. We thank Alek Bolte and Tina Trautmann for their help with the HPAEC-PAD and POC measurements.

## 6. Author contributions

N.S. designed and conducted experiments. S.V.-M. and M.S.-J. took environmental samples. J.-H.H. supervised the project. All authors edited the paper.

