## Supplemental Information for "Biocatalytic quantification of α-glucan in particulate marine organic matter"

| Polysaccharide | Source | Monosaccharide(s) | Glycosidic Linkages |
|---|---|---|---|
| Starch* | Corn | Glucose | α-1,4, α-1,6 |
| Amylopectin* | Corn | Glucose | α-1,4, α-1,6 |
| Amylose* | Potato | Glucose | α-1,4 |
| Glycogen* | Oyster | Glucose | α-1,4, α-1,6 |
| Pullulan† | *Pullularia pullulans* | Glucose | α-1,4, α-1,6 |
| Laminarin* | *Laminaria digitata* | Glucose | β-1,3, β-1,6 |
| Laminarin‡ | *Eisenia bicyclis* | Glucose | β-1,3, β-1,6 |
| Lichenan† | Icelandic moss | Glucose | β-1,3, β-1,4 |
| β-Mannan† | Ivory nut | Mannose | β-1,4 |
| β-Glucan† | Barley | Glucose | β-1,3, β-1,4 |
| Polygalacturonic acid† | Citrus pectin | Galacturonic acid | α-1,4 |
| Xyloglucan† | Tamarind | Xylose, Glucose, Galactose, Arabinose | β-1,4, α-1,6, β-1,6 |
| Carboxymethyl cellulose† | Cellulose | Glucose | β-1,4 |
| α-Mannan* | Yeast | Mannose | α-1,6, α-1,2, α-1,3 |

*Sigma-Aldrich, †Megazyme, ‡Carbosynth

Table S1: Polysaccharide standards used for the results depicted in Figures 1, 2, S2, S3

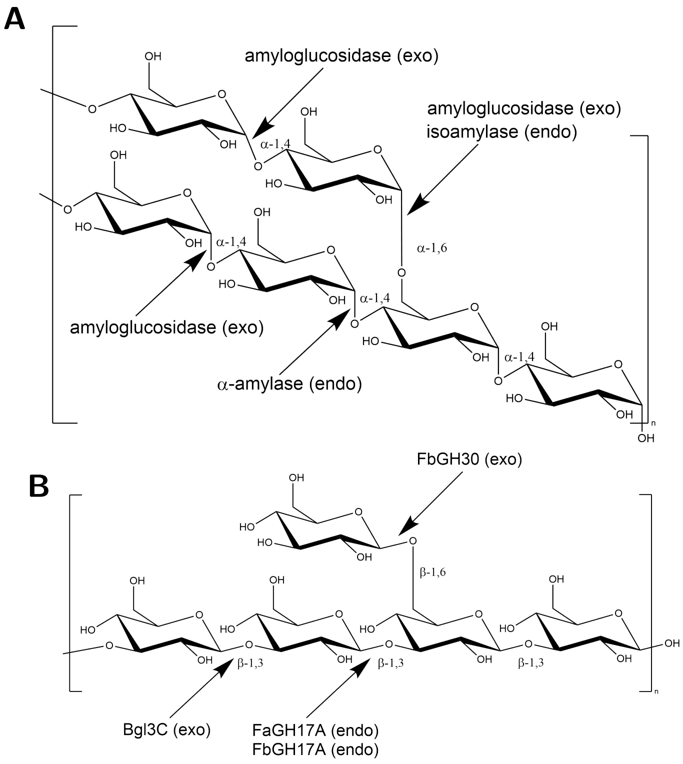


Figure S1: **Partial structures of $\alpha$- and $\beta$-glucans and sites of glycoside hydrolases activities:** (A) A branched $\alpha$-glucan structure as in amylopectin or glycogen with an $\alpha$-1,4-linked $\mathrm{D}$-glucose linear chains and $\alpha$-1,6 branches. The GH13 endo-enzymes $\alpha$-amylase and isoamylase hydrolyze $\alpha$-1,4 linkeages and branching $\alpha$-1,6 linkeages respectively. The exo-acting enzyme amyloglucosidase (GH15) is active on terminal $\alpha$-1,4 or $\alpha$-1,6 glycosidic bonds. (B) The $\beta$-glucan laminarin consists of a $\beta$-1,3-$\mathrm{D}$-glucose polysaccharide main chain with $\beta$-1,6 linked single residue $\mathrm{D}$-glucose side chains. FbGH30 is an exo-acting $\beta$-1,6 glucanase of the GH30 family hydrolysing the branching glucose monomers attached to the laminarin main chain. FaGH17A and FbGH17A are two endo-acting $\beta$-1,3-glucanases of the GH family 17, which act within the $\beta$-1,3-linked laminarin backbone (Unfried et al., 2018; Becker et al., 2017). Bgl3C further hydrolyzes the $\beta$-1,3-$\mathrm{D}$-linked oligos into glucose in an exo-acting manner (Nelson et al., 2017).

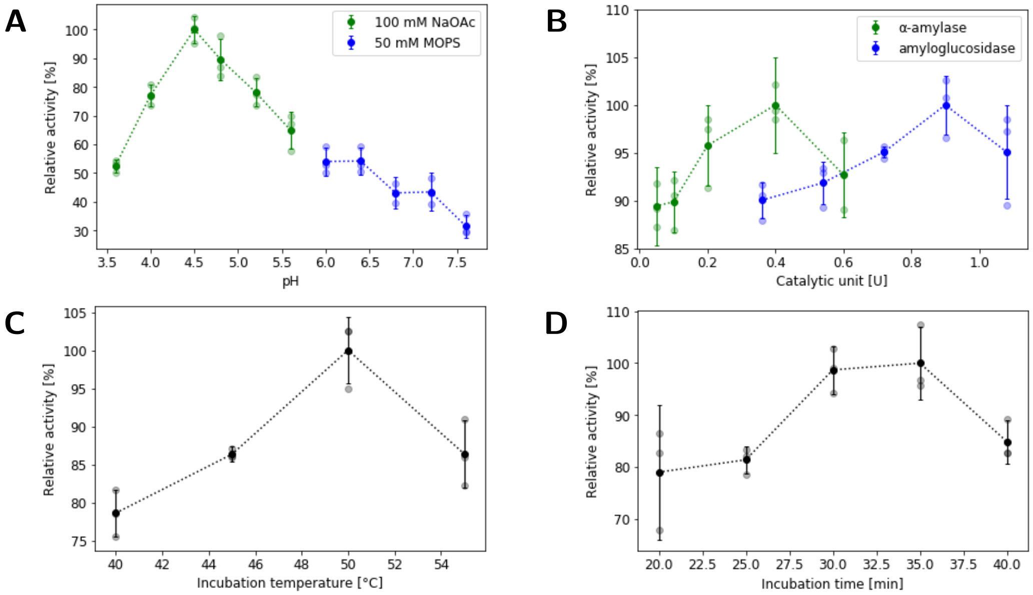


Figure S2: **Optimal conditions for enzymatic $\alpha$-glucan hydrolysis:** Enzymatic activity rates were measured using the PAHBAH reducing sugar assay with 100 µg/mL starch. The greatest activity observed was set as the 100 % reference value for each parameter tested. (A) Comparison of different buffer and pH conditions. (B) Comparison of different enzyme concentrations per sample. (C) Comparison of different incubation temperatures. (D) Comparison of different incubation times.

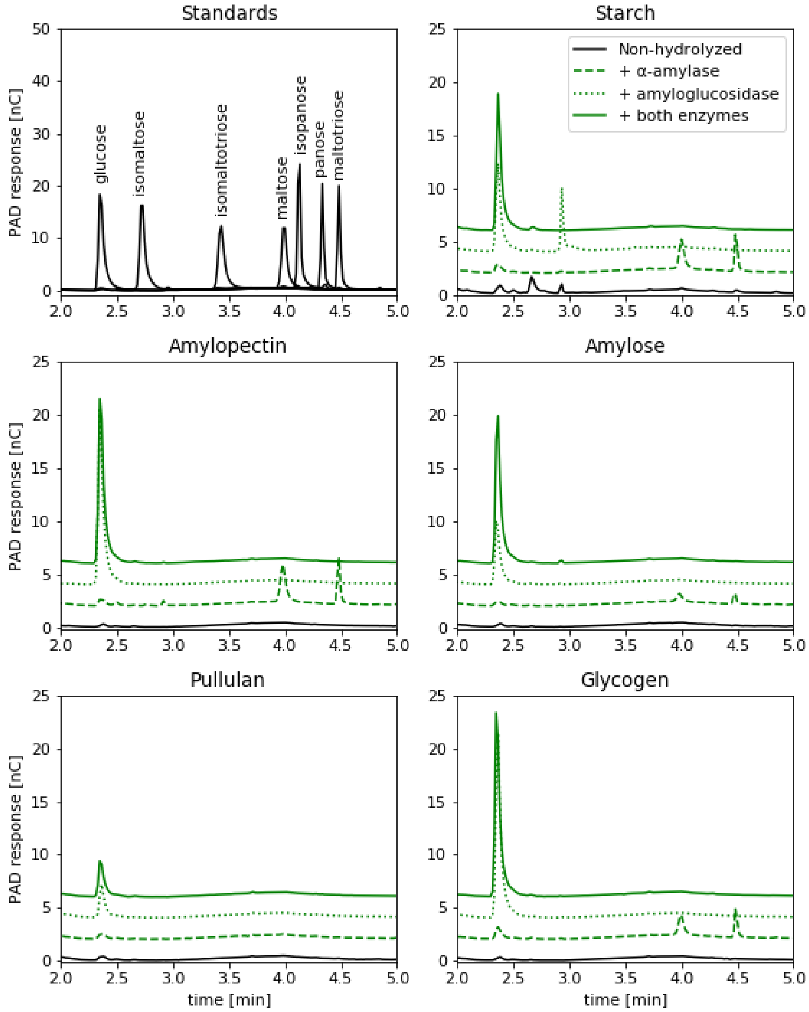


Figure S3: **Enzymatic hydrolysis assays for $\alpha$-glucans produce glucose:** HPAEC-PAD chromatograms (PA200 column) of (top left) maltose standards and glucose, $\alpha$-glucans with and without enzymatic treatment: (top left) starch, (middle left) amylopectin, (middle right) amylose, (bottom left) pullulan and (bottom right) glycogen. Chromatograms of enzymatically hydrolyzed $\alpha$-glucans were moved by 2, 4 and 6 to improve clarity.

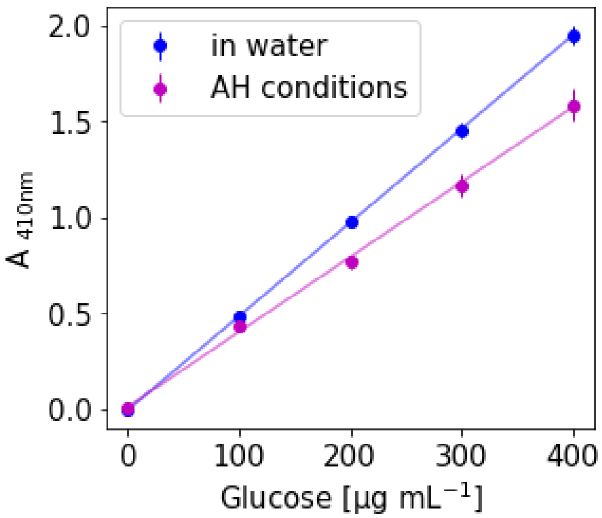


Figure S4: **PAHBAH signal of glucose:** Different concentrations of glucose either directly dissolved in water or subjected to AH conditions.

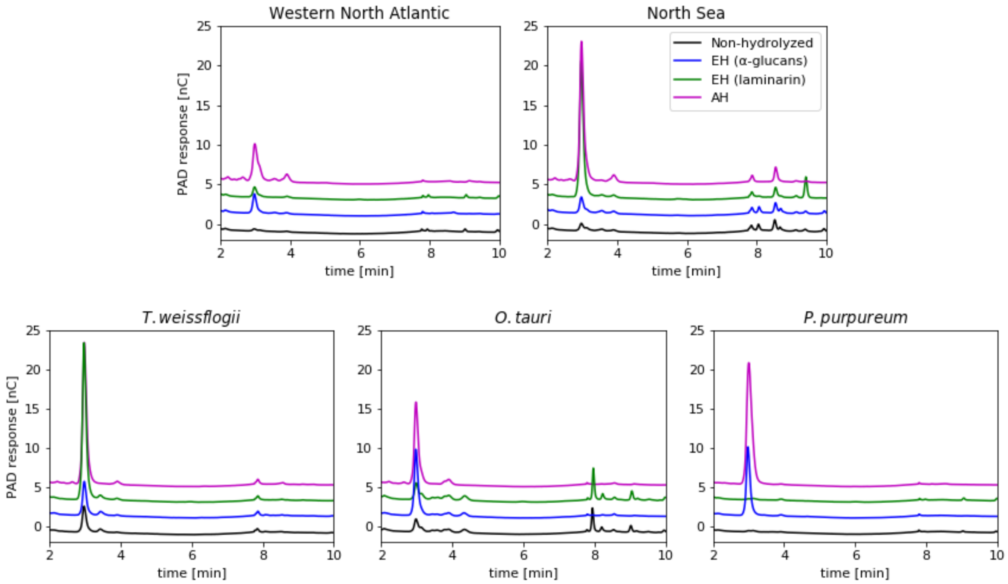


Figure S5: **Enzymatic hydrolysis assays for aqueous extracts produce glucose:** HPAEC-PAD chromatograms (PA100 column) of extracts from (top) environmental POM samples, (bottom) algal cultures. Chromatograms of hydrolyzed samples were moved by 2, 4 and 6 to improve clarity.
